# Meta-analysis reveals host-dependent nitrogen recycling as a mechanism of symbiont control in *Aiptasia*

**DOI:** 10.1101/269183

**Authors:** Guoxin Cui, Yi Jin Liew, Yong Li, Najeh Kharbatia, Noura I Zahran, Abdul-Hamid Emwas, Victor M Eguiluz, Manuel Aranda

**Author notes:** Correspondence to Manuel Aranda.

## Abstract

The metabolic symbiosis with photosynthetic algae of the genus *Symbiodinium* allows corals to thrive in the oligotrophic environments of tropical seas. Many aspects of this relationship have been investigated using transcriptomic analyses in the emerging model organism *Aiptasia*. However, previous studies identified thousands of putatively symbiosis-related genes, making it difficult to disentangle symbiosis-induced responses from undesired experimental parameters. Using a meta-analysis approach, we identified a core set of 731 high-confidence symbiosis-associated genes that reveal host-dependent recycling of waste ammonium and amino acid synthesis as central processes in this relationship. Combining transcriptomic and metabolomic analyses, we show that symbiont-derived carbon enables host recycling of ammonium into nonessential amino acids. We propose that this provides a regulatory mechanism to control symbiont growth through a carbon-dependent negative feedback of nitrogen availability to the symbiont. The dependence of this mechanism on symbiont-derived carbon highlights the susceptibility of this symbiosis to changes in carbon translocation, as imposed by environmental stress.

## Introduction

The symbiotic relationship between photosynthetic dinoflagellates of the genus *Symbiodinium* and corals is the foundation of the coral reef ecosystem. This metabolic symbiosis is thought to enable corals to thrive in the oligotrophic environment of tropical oceans by allowing efficient recycling of nitrogenous waste products in return for photosynthates from the symbionts^1^. Despite the importance of this symbiotic relationship, research has been hampered by the general difficulties associated with the maintenance of corals, their slow growth rates, and the infeasibility of maintaining them in an aposymbiotic state^2^.

To overcome these disadvantages, the sea anemone *Aiptasia* (sensu *Exaiptasia pallida*^3^) has emerged as a powerful model system in the study of cnidarian-*Symbiodinium* symbiosis. *Aiptasia* belongs to the same class (Anthozoa) as corals, and similarly establishes a symbiotic relationship with *Symbiodinium*^4^. In contrast to corals, it can be easily maintained and effectively manipulated under common laboratory conditions. Its rapid asexual reproduction provides relatively large amounts of experimental material for high-throughput studies^5^, while sexual reproduction can be induced efficiently under well-designed conditions^6^. More importantly for symbiosis-related studies, *Aiptasia* can be maintained in an unstressed, aposymbiotic state as long as it is fed regularly^7, 8^. It can also be re-infected with a variety of *Symbiodinium* strains^9^, which allows for comparative studies analyzing the effects of different symbionts on the host. The use of *Aiptasia* as a model organism has advanced our understanding of the metabolic aspects of symbiosis, in particular the identification of glucose as the main metabolite transferred from symbiont to host^10^. However, the molecular mechanisms underlying host-symbiont metabolic interactions are still largely unknown. Particularly the role of nitrogen recycling from waste ammonium is still debated. While it is generally assumed that ammonium assimilation is predominantly performed by the symbiont, some studies show that symbiont-growth is nitrogen limited *in hospite*^11, 12, 13, 14^, suggesting that the host might be able to control nitrogen availability. Consequently, it has been proposed that recycling of ammonium waste by the host might serve as a mechanism to control symbiont densities^15, 16^.

Many genomic, transcriptomic, and proteomic studies have been conducted on the topic of cnidarian-*Symbiodinium* symbiosis in the last two decades to unravel the molecular underpinnings of this relationship ^17, 18, 19, 20, 21, 22, 23^. Due to technical limitations, most of these studies did not have the sensitivity required to detect extensive changes of symbiosis-associated genes. However, these limitations are gradually being overcome by next-generation sequencing techniques. The first *Aiptasia*-centered whole transcriptome comparison between symbiotic and aposymbiotic animals was performed by Lehnert et al.^24^. Since then, multiple studies on the *Aiptasia-Symbiodinium* symbiosis have explored different aspects of this relationship and raised several interesting hypotheses^2, 25^. Despite this increasing wealth of information, our knowledge of underlying key genes associated with this relationship is still limited. While transcriptomic studies have provided valuable information, the resulting lists of putative candidate genes contain thousands of genes, making it difficult to disentangle true symbiosis-related signals from other experimental and technical factors. Furthermore, it was difficult to contrast results across studies due to the lack of a reference genome when most of the studies were carried out. The recent availability of the *Aiptasia* genome^2^ provides a set of high-quality gene models as a reference for transcriptomic analyses. RNA-Seq data can now be mapped directly to these gene models for quantification, thus allowing the comparison of results across different studies.

Here, we carried out a meta-analysis of four RNA-Seq datasets comparing expression differences between symbiotic and aposymbiotic *Aiptasia* (strain CC7) in order to discern sources of technical errors and experimental variations, and to identify a core set of genes and pathways involved in symbiosis establishment and maintenance.

## Results

We conducted our meta-analysis on 3 previous RNA-Seq studies that generated 4 separate datasets, encompassing 17 biological replicates per symbiosis state (i.e., aposymbiotic and symbiotic)^2, 24, 26^.

### Batch effects

In this study, we focused solely on the annotated genes of the previously published *Aiptasia* genome^2^. To investigate the relationship between samples from different studies, we first performed a principal component analysis (PCA) and a rank correlation analysis (RCA) on inter-sample normalized transcripts per million (TPM) values. Both the PCA (Fig. 1A) and RCA (Fig. 1B) showed clear grouping of samples by experiment rather than symbiotic state. However, PCA performed on samples from individual studies showed a clear separation of the samples by symbiotic condition (Fig. S1). This indicates that technical and/or experimental batch effects from each study exert stronger effects on gene expression profiles than the actual symbiotic state of the animals.

**FIG 1.**
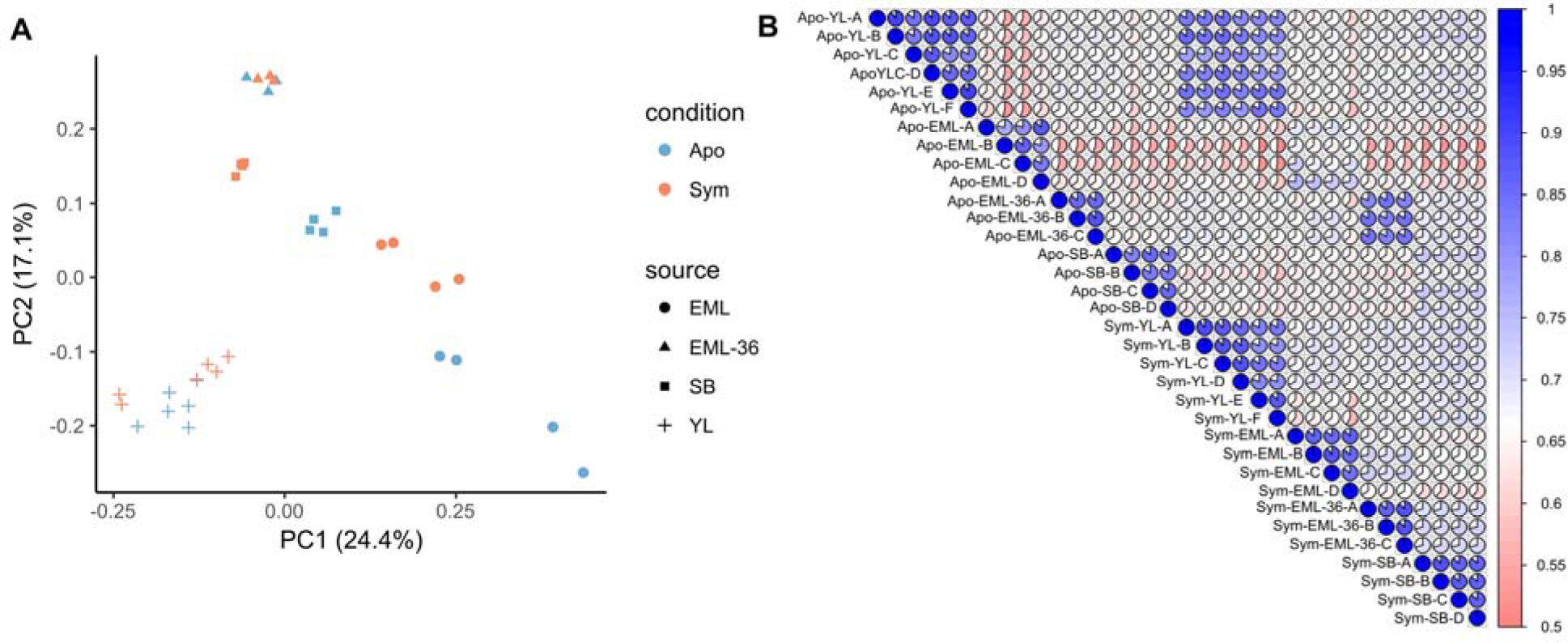
Relationship between samples from different studies. (A) Principal component analysis of samples across all four studies. The symbiotic state (condition) of the animals was indicated by the color of the points, while the source studies were represented as different shapes. (B) Kendall rank correlation of all samples, with high-correlation as blue, and low-correlation as red. The pie chart in each cell also indicates the correlation of the two samples from the corresponding row and column. In both figures, Apo and Sym represent the symbiotic state of the anemones: aposymbiotic and symbiotic, respectively. YL, SB, EML, and EML-36 are the initials of the first authors whose papers we obtained the RNA-Seq data (i.e. Yong Li^26^, Sebastian Baumgarten^2^, and Erik M. Lehnert^24^, respectively).

### Differential expression analyses

Although the four datasets were distinct, there was still a clear separation of symbiotic and aposymbiotic replicates within each of the datasets. We hypothesized that this separation was due to the differential expression of core genes involved in symbiosis initiation and/or maintenance. To identify these genes, we performed four independent differential expression analyses using the exact same pipeline and parameters. These analyses identified between 2,398 to 11,959 differentially expressed genes (DEGs), corresponding to ~10-50% of all expressed genes in the respective studies (Table 1). Surprisingly, the overlap between these lists of DEGs was poor despite the large number of DEGs identified in the individual analyses: only 300 genes were consistently differentially expressed across all four studies. Out of these 300 genes, 166 were upregulated in symbiotic anemones in all comparisons, while 134 were found to be downregulated in symbiotic animals, relative to aposymbiotic controls (Table 1). Paradoxically, we also found 93 genes of 393 genes (23.7%) that were differentially expressed in all studies, but in different directions. At this point, we sought a better technique to identify the core genes involved in symbiosis.

**TABLE 1.** Number of differentially expressed genes in different analyses. “Upregulated” and “downregulated” refers to the number of genes that are expressed at higher levels and lower levels respectively in symbiotic *Aiptasia*, relative to aposymbiotic ones.

| Study | Expressed | DEGs | Upregulated | Downregulated |
| --- | --- | --- | --- | --- |
| YL | 27,684 | 3,058 | 1,552 | 1,506 |
| SB | 24,013 | 11,959 | 6,072 | 5,887 |
| EML | 24,511 | 9,613 | 4,758 | 4,855 |
| EML-36 | 24,246 | 2,398 | 1,241 | 1,157 |
| Overlap | 22,394 | 393 | 166 | 134 |
| Meta-analysis | 25,857 | 731 | 366 | 365 |

### Performing a meta-analysis across four datasets

To obtain a more robust set of core genes involved in symbiosis, we performed a meta-analysis with random effects across the four independent differential gene expression analyses (Table S1). Using this approach, we identified 731 genes that exhibited a more consistent response to symbiosis.

To assess the robustness of these genes, we carried out a principal variance component analysis (PVCA)^27^ to detect the connections between the expression profiles and the different experimental parameters used in each study (Fig. 2, Table S2). For the four individual studies, we found that the symbiotic state of the anemones accounts for a relatively small fraction (6.5% in raw data, 8.4% in normalized data) of the observed variance. Most of the variance was introduced by differences in feeding frequency, days between feeding and sampling, water, light intensity, and temperature. We further noticed that a large proportion of the variance across these four datasets remained unaccountable, suggesting that technical variability, e.g. RNA extraction, library preparation and sequencing, also introduces substantial unwanted heterogeneity to gene expression profiles. When the PVCA was similarly applied to the 731 genes identified through our meta-analysis, we observed that these genes had a significantly enhanced association with symbiosis. Symbiosis state accounted for 46.6% of the expression variance observed in these genes (Fig 2).

We noticed that smaller gene lists tended to have variances that were better explained by symbiosis state, exemplified by DEG_YL and DEG_EML-36 having better association with symbiosis than DEG_SB and DEG_EML. Thus, one could argue that the meta-analysis merely achieved better association with symbiosis as it had the fewest genes of interest. To assess this confounding factor, we performed PVCA on a set of randomly picked 731 genes from DEG_YL. This was repeated 10,000 times (i.e., a Monte-Carlo approach), and for other DEG lists (DEG_SB, DEG_EML and DEG_EML-36). These simulations allowed us to estimate that the likelihood of our meta-analysis producing the observed 46.6% by random chance was *p* < 10^−4^ (0 of 40,000 trials had symbiosis state accounting for > 46.6% of the variance).

**FIG 2.**
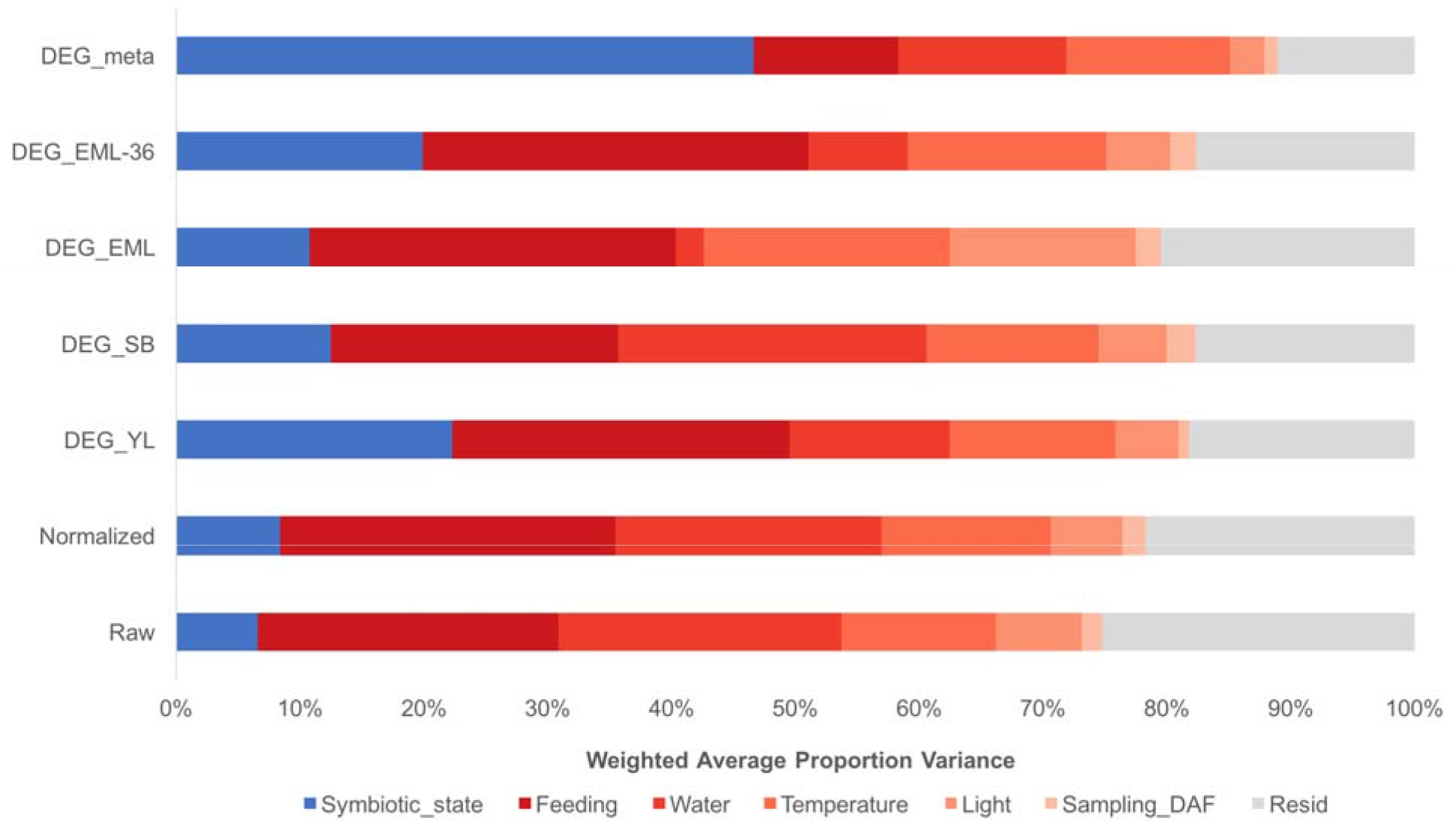
Principal variance component analysis of DEGs from different analyses. The contribution of each factor to the overall variance in each analysis was estimated by PVCA. The variance explained by symbiotic state (blue) is highest in the set of DEGs from the meta-analysis (DEG_meta); the combined variation attributable to experimental factors (red) is lowest in DEG_meta as well. Unresolved variance is in gray. DEG_YL, _SB, _EML and _EML-36 represents the set of differentially expressed genes identified in four independent differential analyses. Raw and Normalized are the combined raw and inter-sample normalized expression data across all *Aiptasia* genes, showing that < 10% of the variation in overall gene expression can be attributed to symbiotic state. DAF: days after feeding.

### Functional interpretation

To assess the impact of the previously identified experiment-specific biases, we conducted GO and KEGG pathway enrichment analyses on the DEGs identified using the four independent differential gene expression analyses, respectively. Across the analyses of four independent experiments, 283-645 GO terms and 9-55 KEGG pathways were enriched. However, the functional overlap across all studies was poor: a large proportion of the putatively enriched terms were only identified in a single dataset (~75% in GO, and ~65% in KEGG) (Fig. S2). Compared to these independent analyses, the GO and KEGG pathway enrichment of the 731 symbiosis-associated core genes contained fewer significant GO terms (204), but comparatively more significantly enriched KEGG pathways (31). Many of the enriched GO terms and KEGG pathways, as well as their associated genes, fit well with processes previously reported to be involved in symbiosis, including symbiont recognition and the establishment of symbiosis, host tolerance of symbiont, and nutrient exchange between partners and host metabolism which are discussed separately (Supplementary Information S1). However, our analysis also identified several symbiosis-related processes that were previously overlooked; of these processes, pathways associated with amino acid metabolism exhibited the most extensive changes in response to symbiosis.

### Extensive changes of amino acid metabolism in response to symbiosis

Amino acid and protein metabolism represented a major symbiosis-related aspect in our meta-analysis. 9 of 31 enriched KEGG pathways and 18 of 125 enriched biological process GO terms were associated with amino acid and/or protein metabolism (Fig. 3). A total of 97 DEGs were involved in these processes, of which 43 were upregulated in symbiotic animals. Interestingly, the DEGs involved in most of the enriched biological processes exhibited consistent expression changes (Fig. 3A), i.e. the genes associated with the corresponding process were either exclusively upregulated or downregulated.

**FIG 3.**
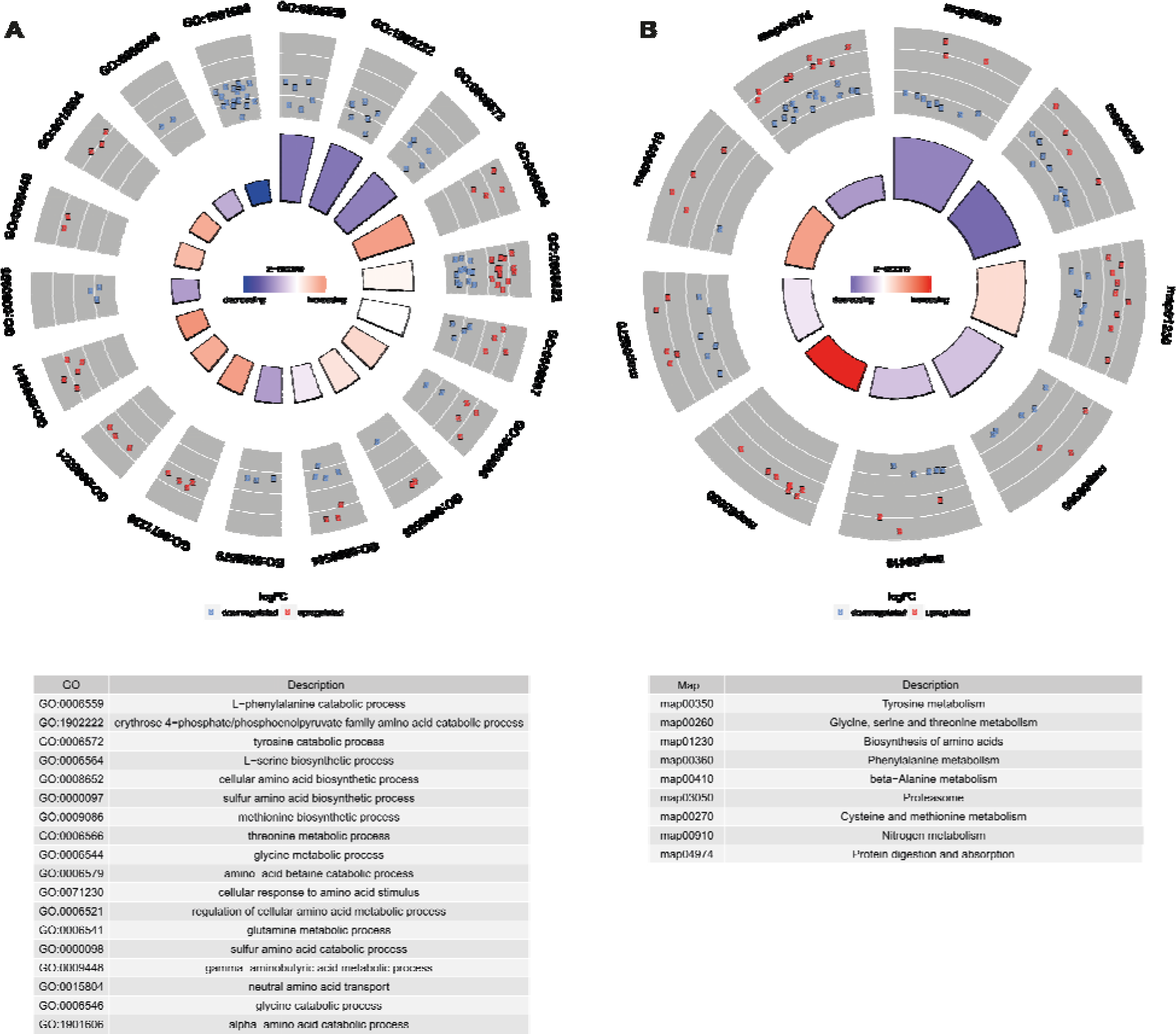
Amino acid metabolism biological processes (A) and pathways (B) enriched with DEGs identified in meta-analysis. For the two Circos plots, the height of each bar in the inner circle indicates statistical significance of the enriched GO terms (A) and KEGG pathways (B), while color of the bars represents the overall regulation effect of each process. The outer circle shows the differential expression of genes associated with each process, where red and blue represent upregulation and downregulation in symbiotic anemones, respectively. The table describes the annotation of each term or pathway.

Further integration of these enriched biological processes and pathways revealed an amino acid metabolism hub in *Aiptasia-Symbiodinium* symbiosis (Fig. 4). We observed that genes catalyzing glycine/serine biosynthesis from food-derived choline were systematically downregulated in symbiotic anemones. In contrast, the genes involved in *de novo* serine biosynthesis from 3-phosphoglycerate, one of the glycolysis intermediates, and glutamine/glutamate metabolism were generally upregulated (Fig. 4A). The resulting change in amino acid synthesis pathways suggested that symbiotic hosts utilize glucose and waste ammonium to synthesize serine and glycine, which are both main precursors for many other amino acids (Supplementary Information S1). Based on these findings, we hypothesized that the host uses symbiont-derived glucose to assimilate waste ammonium to produce amino acids.

To test this hypothesis, we further investigated metabolomes of symbiotic and aposymbiotic anemones using nuclear magnetic resonance (NMR) spectroscopy. Three metabolites in the *de novo* serine biosynthesis pathway were highly abundant in symbiotic *Aiptasia* (two of them significantly so, *p* < 0.05), while five out of the six intermediates in the alternative glycine/serine biosynthesis pathway using food-derived choline were significantly enriched in aposymbiotic anemones as predicted (Fig. 4B). However, as glucose produces multiple peaks in the ^1^H NMR spectrum, and most of these peaks overlap with many other potential metabolites in both symbiotic and aposymbiotic anemones, it was not possible to precisely determine glucose concentrations via NMR. Consequently, we performed ^13^C bicarbonate labeling experiments and compared metabolite profiles of symbiotic and aposymbiotic anemones using gas chromatography-mass spectrometry (GC-MS), in order to test if the glucose is indeed provided by the symbiont and if the downstream usage of symbiont derived organic carbon is in the host. Our experiments confirmed that symbionts provide large amounts of ^13^C-labeled glucose to the host (Fig. S3) and that the ^13^C-labeling was significantly enriched in many amino acids and their precursors in symbiotic anemones compared to aposymbiotic ones (Table S2). Moreover, metabolite set enrichment analysis indicates that these ^13^C-enriched are associated mainly with several amino acid metabolism pathways (Fig. S4), which is consistent with the enrichment analysis of 731 differentially expressed genes. For the amino acids with good abundance in both symbiotic and aposymbiotic animals, we examined the proportion of ^13^C in each of them, respectively. Interestingly, we observed relatively stable increases (~1.5-fold) of ^13^C levels in symbiotic animals compared with aposymbiotic ones (Fig. 4C). This constant increase may indicate there is a unique carbon source (photosynthesis-produced glucose) rather than multiple sources (glucose and symbiont-derived amino acids) in host amino acid biosynthesis.

**FIG 4.**
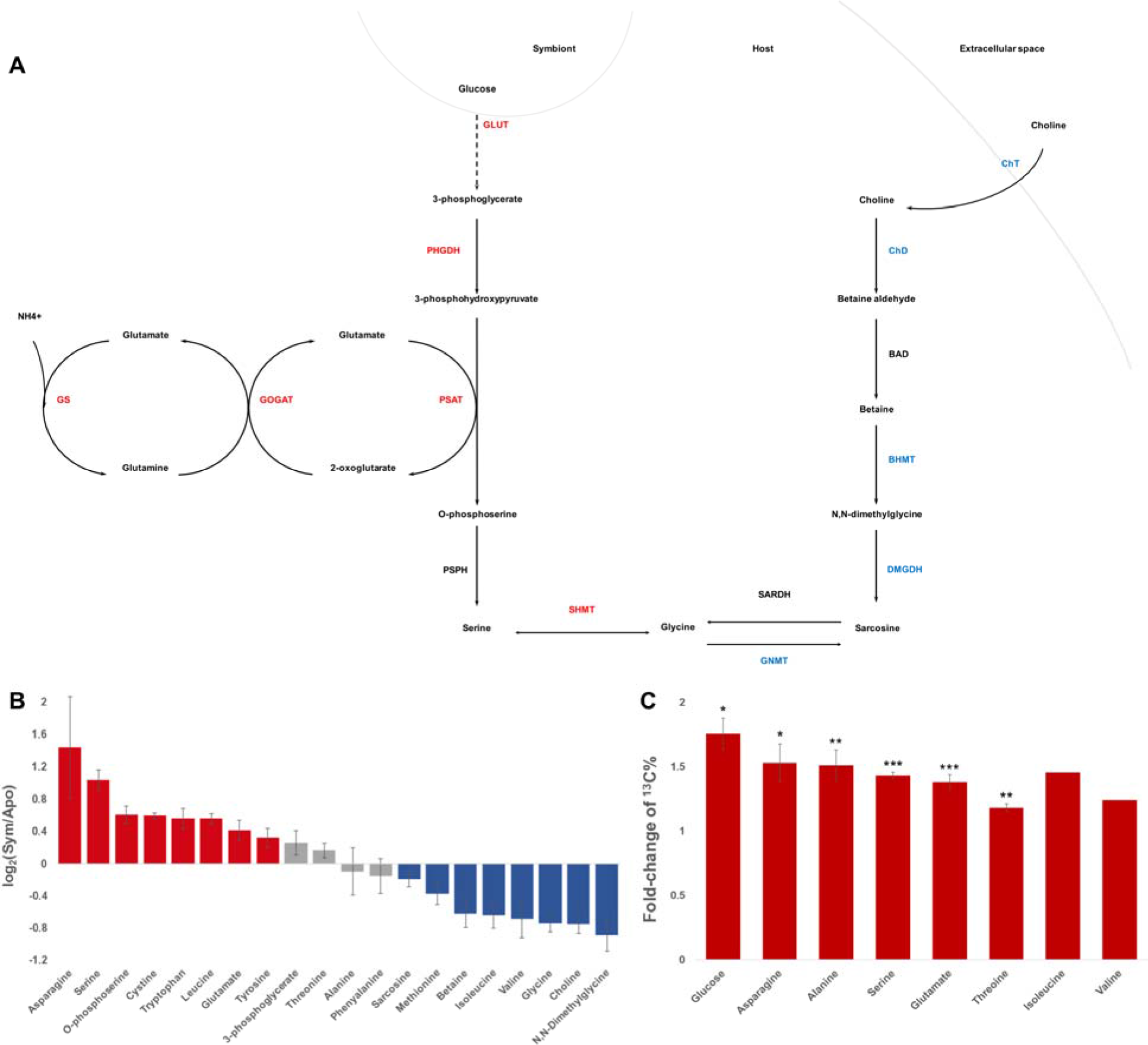
Amino acid metabolism in *Aiptasia-Symbiodinium* symbiosis. (A) Serine biosynthesis in *Aiptasia* with different symbiotic states. The pathway on the left indicates *de novo* serine biosynthesis from symbiont-produced glucose, while the right part represents glycine/serine biosynthesis from food-derived choline. Enzyme names are colored to indicate differential expression of the corresponding genes, where red and blue mean upregulation and downregulation in symbiotic anemones, respectively. (B) Metabolite abundance changes in response to symbiosis. Color represent abundance changes, with red for significant increases in symbiotic anemones, blue for significant increase in aposymbiotic animals, and gray for non-significant changes. (C) Increasing ^13^C proportion of glucose and amino acids in symbiotic *Aiptasia*. Asterisks denote statistical significance of the changes (two-tail *t*test: * *p* < 0.05, ** *p* < 0.01, *** *p* < 0.001). Statistical testing of isoleucine and valine was not possible as they were detected in only one aposymbiotic replicate with reasonable concentration. Error bar represents standard error of the mean.

## Discussion

Batch effects are known to introduce strong variation in high throughput sequencing studies^28, 29^. However, this is often overlooked in transcriptomic studies, and especially so in non-model organisms. Our analysis of RNA-Seq data from four independent experiments analyzing transcriptional changes between symbiotic and aposymbiotic *Aiptasia* highlighted that batch effects are indeed pervasive in published data, even among studies using the same genotype (clonal strain CC7). Analyses of the combined dataset from all four experiments showed clear grouping of samples by experiment rather than treatment. However, when each experiment was analyzed independently, replicates separated by symbiotic states as expected. Interestingly, we found that the observed batch effects were not restricted to technical biases. Our analyses showed that the specific experimental setups in each study were a greater source of variance than the symbiosis state, which was the actual factor of interest in these studies. More importantly, we found that genes closely related to the processes involved in symbiosis, such as nutrient exchanges, may also respond significantly to various parameters of culture conditions, such as the feeding frequency, days between sampling and feeding, water, light intensity, and the temperature. Without careful design, such factors may exert effects on gene expression that mask the changes specific to the treatment of interest (symbiotic state).

Based on our findings, we suggest two potential venues to reduce the high signal-to-noise ratio in differential expression studies. Firstly, future transcriptomic efforts should take extreme care to standardize all experimental conditions save for the one under study. For example, culture conditions should be identical, treatments should be performed on multiple independent batches, RNA extractions and library preparation should be carried out on all samples simultaneously. The prepared libraries should also be sequenced in the same run to further minimize technical variations. Secondly, one should not dogmatically adhere to the convention of using *p* = 0.05 as the cutoff for statistical significance. If a study considers one in every three genes as significantly differentially expressed, to a careful reader, the proclaimed significance of those genes is diminished. As the number of DEGs increase, the rate of type I errors would also increase, which would make the discovery of meaningful biological processes more difficult.

From the functional interpretation of DEGs associated with enriched GO terms and KEGG pathways, we found that many processes in the host were significantly induced or suppressed in response to symbiosis. One of the key features that has been overlooked in previous studies is the switch of serine biosynthesis pathways in *Aiptasia* in response to symbiosis.

The downregulation of choline transport indicates a decrease of the host’s demand on dietary choline during symbiosis. Correspondingly, genes involved in the downstream conversion of choline to betaine and the production of glycine from betaine are also downregulated. The decrease of glycine caused by this downregulation is likely compensated by the metabolism of serine, which can be achieved by the observed upregulation of serine hydroxymethyltransferase (SHMT, AIPGENE4781), which catalyzes the interconversions between glycine and serine. Interestingly, our results suggest that serine is one of the key components in the amino acid interconversions, as the genes involved in its *de novo* biosynthesis from 3-phosphoglycerate (one of the intermediates of glycolysis) were consistently upregulated. The conversion from glutamate to 2-oxoglutarate, catalyzed by the upregulated phosphoserine aminotransferase (PSAT, AIPGENE17104), may serve as the main reaction to provide amino groups for the biosynthesis of amino acids. Since 2-oxoglutarate is also one of the intermediates in the citrate acid cycle, an increase of glucose provided by the symbionts may also increase the overall activity of the cycle, hence raising the relative abundance of 2-oxoglutarate in symbiotic animals. High levels of 2-oxoglutarate have been reported to induce ammonium assimilation through glutamine synthetase / glutamate synthase cycle^30^. Consistent with this finding, we observe all the genes involved in this pathway to be upregulated in symbiotic anemones.

Metabolomic analyses of symbiotic and aposymbiotic anemones confirm the predictions derived from our transcriptomic meta-analysis. Most of the intermediates in the *de novo* serine biosynthesis using symbiont-derived glucose were highly enriched in symbiotic anemones and showed increased ^13^C-labeling. whereas many of the metabolites from choline-betaine-glycine-serine conversion have decreased abundance in symbiotic animals. Furthermore, we also identified many other amino acids showing significantly increased abundance and ^13^C-labeling signals, suggesting that serine may serves as metabolic intermediate for the production of other amino acids. Taken together, these results highlight that symbiont-derived glucose fuels ammonium assimilation and amino acid production in the host and that serine biosynthesis acts as a main metabolic hub in symbiotic hosts.

The strong shifts in host amino acids metabolic pathways induced by symbiont-provided glucose described here indicate the major nitrogen and carbon sources of the anemone host, and their interactions in the *Alptasla-Symblodlnlum* symbiosis. The catabolism of glucose through pathways such as glycolysis, pentose phosphate pathway, and citric acid cycle, not only generates more energy (in forms of ATP, NADH, and NADPH), which is critical to ammonium assimilation, but also produces more intermediate metabolites that can serve as carbon backbones in many biosynthetic pathways such as amino acid synthesis. Our findings thus highlight nitrogen conservation, i.e. the host driven assimilation of waste ammonium using symbiont-derived carbon, as a central mechanism of the cnidarian-algal endosymbiosis^16^. This metabolic interaction might serve as a self-regulating mechanism for the host to control symbiont density through the regulation of nitrogen availability^15^ in a carbon dependent manner. This allows for higher nitrogen availability in early stages of infection (few symbionts translocating few carbon) and gradual reduction of nitrogen availability with increasing symbiont densities (many symbionts translocating more carbon). The strict dependence of this mechanism on symbiont-derived carbon highlights the sensitivity of this relationship to changes in carbon translocation as imposed by stress-induced retention of photosynthates by symbionts^31, 32^.

## Materials and Methods

### Data collection and pre-processing

Based on literature review of recently published *Alptasla* genome and transcriptome studies, four datasets generated from three previous publications^2, 24, 26^ were obtained (Table 1). All RNA-Seq experiments were performed on the clonal *Aiptasia* strain (CC7) and sequenced on the same platform (Illumina HiSeq 2000). Three of the datasets contained 101 bp paired-end reads, while the last one contained 36 bp single-end reads. Samples were labeled based on the initials of the first author of published papers and ongoing project. As all raw data from Lehnert et al.^24^ was provided as a monolithic FASTQ file, a custom Python script was written to split the reads into its constituent replicates, as inferred from the FASTQ annotation lines.

**Table 1.**
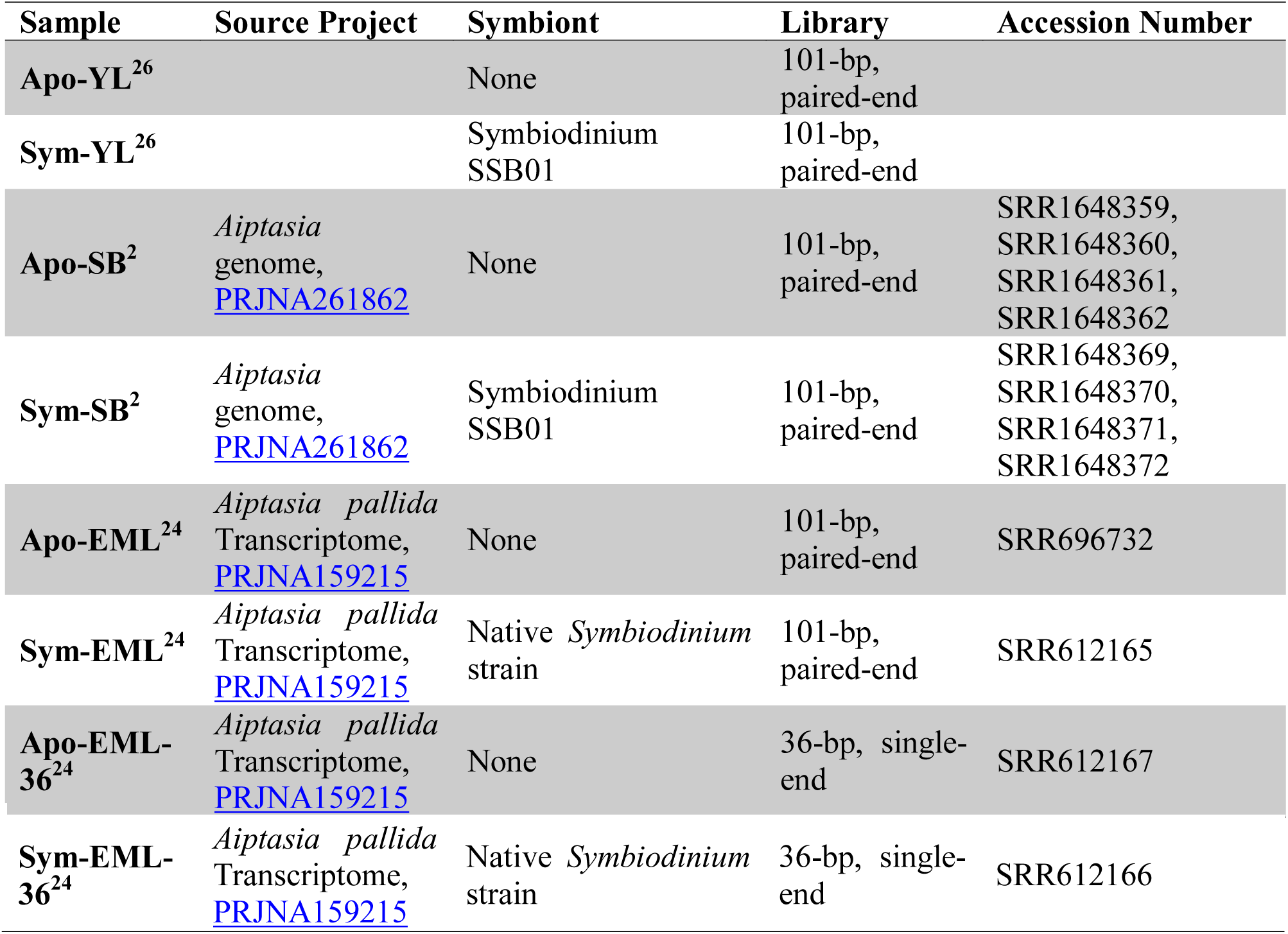
Summary of the NGS data sources used in this study

| Sample | Source Project | Symbiont | Library | Accession Number |
| --- | --- | --- | --- | --- |
| **Apo-YL<sup>26</sup>** |  | None | 101-bp, paired-end |  |
| **Sym-YL<sup>26</sup>** |  | Symbiodinium SSB01 | 101-bp, paired-end |  |
| **Apo-SB<sup>2</sup>** | *Aiptasia* genome, [PRJNA261862](https://www.ncbi.nlm.nih.gov/nuclink/PRJNA261862) | None | 101-bp, paired-end | SRR1648359, SRR1648360, SRR1648361, SRR1648362 |
| **Sym-SB<sup>2</sup>** | *Aiptasia* genome, [PRJNA261862](https://www.ncbi.nlm.nih.gov/nuclink/PRJNA261862) | Symbiodinium SSB01 | 101-bp, paired-end | SRR1648369, SRR1648370, SRR1648371, SRR1648372 |
| **Apo-EML<sup>24</sup>** | *Aiptasia pallida* Transcriptome, [PRJNA159215](https://www.ncbi.nlm.nih.gov/nuclink/PRJNA159215) | None | 101-bp, paired-end | SRR696732 |
| **Sym-EML<sup>24</sup>** | *Aiptasia pallida* Transcriptome, [PRJNA159215](https://www.ncbi.nlm.nih.gov/nuclink/PRJNA159215) | Native *Symbiodinium* strain | 101-bp, paired-end | SRR612165 |
| **Apo-EML-36<sup>24</sup>** | *Aiptasia pallida* Transcriptome, [PRJNA159215](https://www.ncbi.nlm.nih.gov/nuclink/PRJNA159215) | None | 36-bp, single-end | SRR612167 |
| **Sym-EML-36<sup>24</sup>** | *Aiptasia pallida*<br>Transcriptome,<br>PRJNA159215 | Native *Symbiodinium*<br>strain | 36-bp, single-end | SRR612166 |

### Identification of DEGs

To avoid biases stemming from the use of disparate bioinformatics tools in calling DEGs, data from the four datasets were processed with identical analytical pipelines.

Gene expressions were quantified (in TPM, transcripts per million) based on the published *Aiptasia* gene models^2^ using kallisto v0.42.4^33^. DEGs were independently identified in the four datasets using sleuth v0.28.0^34^. Genes with corrected *p* values < 0.05 were considered differentially expressed.

To enable direct comparisons of gene expression values between datasets, another normalization with sleuth was carried out on all samples (*n* = 17 aposymbiotic and *n* = 17 symbiotic). Principal component analysis (PCA) and ranked correlation analysis (RCA) were carried out on these normalized expression values to assess the relationship between samples and reproducibility of these studies.

### Profiling sources of batch effects

Principal variance components analysis (PVCA), a technique that was developed to estimate the extent of batch effects in microarray experiments^27^, was used several times in our study. A PVCA was carried out on raw data to estimate the batch effects in the combined dataset and their possible source in the original experimental designs; similarly, the normalized data was also assessed for the reduction of batch effects post-normalization. We also performed PVCA on normalized expression values of the differentially expressed genes (DEG) identified in each independent analysis or the final meta-analysis to detect the robustness of DEG calling.

### Meta-analysis across studies

For every gene with at least two studies with significant differential expression values, a meta-analysis was performed to determine the overall effect size and associated standard error. Effect sizes from each study *i* (represented as *w*_*i*_) were calculated as the natural logarithm of its expression ratio (ln *R*_*i*_), i.e. geometric means of all expression values in the aposymbiotic state divided by the geometric means of all expression values in the symbiotic state. Conveniently, this value is approximately equal to the *β*_*i*_ value provided by sleuth. As sleuth also calculates the standard error of *β*_*i*_, the variance of ln *R*_*i*_ was not calculated via the typical approximation—instead, the variance *ν*_*i*_ was directly calculated as

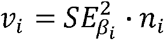

where *n*_*i*_ represents the number of replicates in study *i*.

To combine the studies, a random-effects model was used. While the use of this model is somewhat discouraged for meta-analyses with few studies as it is prone to produce type I errors^35^, we still opted for its use over the fixed-effects model due to the substantial inter-study variation evident in the PCAs performed previously. Also, the type I error rate could be controlled by setting a more conservative *p* threshold, if required.

The DerSimonian and Laird ^36^ method was implemented as described below. Studies with individual effect sizes *m*_*i*_ were weighted (*w*\*) by a combination of the between-study variation (*τ*^2^) and within-study variation (*ν*_*i*_), according to the formula

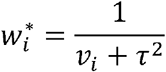

The between-study variation (*τ*^2^) across all *k* studies was calculated as

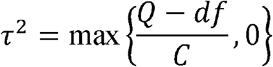

where

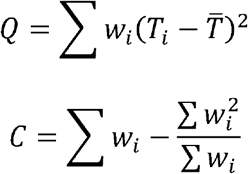

The weighted mean (*m*\*) was calculated as

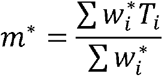

while the standard error of the combined effect was

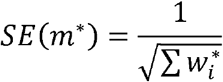

The two-tailed p-value was calculated using

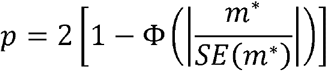

and then subsequently corrected for multiple hypothesis testing with the Benjamini-Hochberg-Yekutieli procedure^37, 38^ using a Python script. Genes with corrected *p* < 0.05 were considered differentially expressed. For transparency, calculations for all equations were implemented manually in Microsoft Excel (Table S3) following established guidelines^39^.

### Functional interpretation of DEGs

Gene ontology (GO) and KEGG pathway enrichment analyses were both conducted on five DEG lists: one each from the four independent datasets, and one from the results of the meta-analysis.

Identification of enriched GO terms were conducted using topGO^40^ by a self-developed R script (https://github.com/lyijin/topGO_pipeline). A GO term was considered enriched only when its *p* value was less than 0.05.

KEGG pathway enrichment analyses were performed using Fisher’s exact and subsequent multiple testing correction via false discovery rate (FDR) estimation. A KEGG pathway was deemed enriched (or depleted) only when its FDR less than 0.05. The results of enrichment analyses were visualized using GOplot^41^.

### Metabolomic profiles of symbiotic and aposymbiotic anemones

*Aiptasla* strain CC7 was bleached and re-infected with a compatible strain of *Symblodlnlum* SSB01 as previously reported^2^. All the symbiotic and aposymbiotic anemones were maintained in the laboratory in autoclaved seawater (ASW) at 25 °C in 12-hour light/12-hour dark cycle with light intensity of ~30 μmol photons m^−2^s^−1^ for over three years. Anemones were fed three times a week with freshly hatched *Artemia* nauplii, and water change was done on the day after feeding.

Anemones were rinsed extensively to remove any external contaminations, and starved for two days in ASW and transferred into ASW with 10 mM ^13^C-labelled sodium bicarbonate (Sigma-Aldrich, USA) for another two days before sampling. The four-day starvation period ensured all *Artemia* had been digested and consumed, hence there was no contamination from *Artemia* in the samples for NMR and GC-MS. The samples were drained completely on clean tissues to remove any water on surface, then snap frozen in liquid nitrogen to avoid any further metabolite changes in downstream processing.

To compare metabolomic profiles of anemones at different symbiotic states, four replicates of each state (*n* = 30 individuals per replicate), were processed for metabolite extraction using a previously reported methanol/chloroform method^42^. The free amino acid-containing methanol phase was dried using CentriVap Complete Vacuum Concentrators (Labconco, USA).

For NMR metabolite profiling, samples were dissolved in 600 μl of deuterated water (D_2_O), and vortexed vigorously for at least 30 seconds. Subsequently, 550 μL of the solution was transferred to 5 mm NMR tubes. NMR spectrum was recorded using 700 MHz AVANCE III NMR spectrometer equipped with Bruker CP TCI multinuclear *CryoProbe* (BrukerBioSpin, Germany). To suppress any residual HDO peak, the ^1^H NMR spectrum were recorded using excitation sculpting pulse sequence (zgesgp) program from Bruker pulse library. To achieve a good signal-to-noise ratio, each spectrum was recorded by collecting 512 scans with a recycle delay time of 5 seconds digitized into 64 K complex data points over a spectral width of 16 ppm. Chemical shifts were adjusted using 3-trimethylsilylpropane-1-sulfonic acid as internal chemical shift reference. Before Fourier transformations, the FID values were multiplied by an exponential function equivalent to a 0.3 Hz line broadening factor. The data was collected and quantified using Bruker Topspin 3.0 software (Bruker BioSpin, Germany), with metabolite-peak assignment using Chenomx NMR Suite v8.3, with an up-to-date reference library (Chenomx Inc., Canada).

For ^13^C-labelling investigation using GC-MS, dried samples were re-dissolved in 50 μl of Methoxamine (MOX) reagent (Pierce, USA) at room temperature and derivatized at 60 °C for one hour. 100 μl of *N,O-bis*-(trimethylsilyl) trifluoroacetamide (BSTFA) was added and incubated at 60 °C for further 30 min. 2 μl of the internal standard solution of fatty acid methyl ester (FAME) were then spiked in each sample and centrifuged for 5 min at 10,000 rpm. 1 μl of the derivatized solution was analyzed using single quadrupole GC-MS system (Agilent 7890 GC/5975C MSD) equipped with EI source at ionization energy of 70 eV. The temperature of the ion source and mass analyzer was set to 230 °C and 150 °C, respectively, and a solvent delay of 9.0 min. The mass analyzer was automatically tuned according to manufacturer’s instructions, and the scan was set from 35 to 700 with scan speed 2 scans/s. Chromatography separation was performed using DB-5MS fused silica capillary column (30m × 0.25 mm I.D., 0.25 μm film thickness; Agilent J&W Scientific, USA), chemically bonded with 5% phenyl 95% methylpolysiloxane cross-linked stationary phase. Helium was used as the carrier gas with constant flow rate of 1.0 ml min^−1^. The initial oven temperature was held at 80◻C for 4 min, then ramped to 300 °C at a rate of 6.0 °C min-1, and held at 300 °C for 10 min. The temperature of the GC inlet port and the transfer line to the MS source was kept at 200 °C and 320 °C, respectively. 1 μl of the derivatized solution of the sample was injected into split/splitless inlet using an auto sampler equipped with 10 μl syringe. The GC inlet was operated under splitless mode. Metabolites in all samples were identified using Automated Mass Spectral Deconvolution and Identification System software (AMDIS) with the NIST special database 14 (National Institute of Standards and Technology, USA). The mass isotopomer distributions (MIDs) of all compounds were detected and their ^13^C-labelling enrichment in symbiotic *Aiptasia* were investigated using MIA^43^. Pathways associated with these ^13^C-enriched metabolites were explored using MetaboAnalyst v3.0^44^.

## Author Contributions

M.A. conceived the project. G.C., Y.J.L., M.A., Y.L., N.I.Z., and V.M.E. collected the RNA-Seq data, performed data analyses, and visualized the results. G.C., N.K., A.E., and M.A. conducted metabolomic experiments and analyzed the data. G.C., Y.J.L., and M.A. wrote the manuscript with method input from other authors. All authors have read and agreed on the final draft.

## Acknowledgements

We would like to thank Jit Ern Chen and Maha J. Cziesielski for valuable comments on our manuscript. This publication is based upon work supported by the King Abdullah University of Science and Technology (KAUST) Office of Sponsored Research (OSR) under Award No. URF/1/2216-01.

